# The Transcriptomics Pain Signature Database

**DOI:** 10.1101/2023.06.16.545337

**Authors:** Sahel Jahangiri Esfahani, Marc Parisien, Calvin Surbey, Luda Diatchenko

**Author notes:** These authors equally contributed to this work (Sahel Jahangiri Esfahani, Marc Parisien).

## Abstract

The availability of convenient tools is critical for the efficient analyses of fast-generated omics-wide-level studies. Here, we describe the creation, characterization, and applications of the Pain Signatures Database (TPSDB), a comprehensive database containing the results of differential gene expression analyses from 338 full transcriptomic datasets for pain-related phenotypes. The database allows searching for a specific gene(s), pathway(s), or SNP(s), or downloading the raw data for hypothesis-free analysis. We took advantage of this unique dataset of multiple pain transcriptomics in several ways. The pathway analyses found the cytokine production regulation and innate immune response the most frequently shared pathways across tissues and conditions. A machine learning-based approach across datasets identified RNA biomarkers for inflammatory and neuropathic pain in rodent dorsal root ganglion (DRG) with high certainty. Finally, functional annotation of pain-related GWAS results demonstrated that differentially expressed genes can be more informative than the general tissue-specific genes from DRG or spinal cord in partitioning heritability analyses.

## INTRODUCTION

The International Association for the Study of Pain defined pain as “an unpleasant sensory and emotional experience associated with, or resembling that associated with actual or potential tissue damage”^1^. Pain is not always indicative of tissue injury and it can persist, even long after a trauma or inflammatory event^2^. The pain that lasts longer than it can serve any protective role is called chronic and is a medical problem itself. Such pain lowers people’s quality of life and increases the associated medical costs^3^. These challenges add to the importance of pain research.

The molecular architecture of pain and pain resolution is started to be understood^4–6^. The identification of such molecular pathways is critical for both biomarker development and drug target discovery, as well as for building a thorough map of the molecular pathophysiology of pain states. Like in other complex diseases, a variety of factors can influence the way we perceive, resolve, and respond to pain. Therefore, multiple omics-wide level studies are needed to capture as many involved genes and pathways as possible. Furthermore, omics-wide studies are hypothesis-free and allow the identification of new genes and pathways. These studies have become feasible through advances in sequencing and other high-throughput techniques^7^.

Since 1996, with public data sharing policies, it has been made possible for researchers to benefit from other researchers’ shared data, leading to noteworthy scientific progress in genetics^8^. Nevertheless, not all individuals can use these genetic data in their publicly available format, and specific data analysis tools and skills are needed to be able to use them directly or convert them into a meaningful format. Also, even for bioinformaticians, it would be laborious to analyze genetic data from multiple sources, even if it is a few datasets.

We thus introduce here the Transcriptomics Pain Signature Database (TPSDB), where we organized publicly available full transcriptomics datasets to conveniently search for the differentially expressed genes (DEGs) for pain-related perturbations. TPSDB makes it possible for users to easily check the trajectory of their genes-of-interest or pathway in various organisms, tissues, time points, sexes, pain phenotypes, and assays, in the context of pain. We first did a thorough characterization of this unique dataset of multiple pain transcriptomics that provided many insights into critical genes and pathways implicated in pain. Furthermore, we demonstrated here the utility of our transcriptomics database for two applications. The first application was the RNA biomarkers identification. Using a machine learning approach, we identified rodents’ dorsal root ganglion (DRG) RNA markers of neuropathic pain (NP), inflammatory pain (IP), and sham in the corresponding pain assays. The second application was the functional analysis of genome-wide association studies (GWAS) results. Here, we performed a partitioned heritability analysis using the genes obtained from the differential gene expression analyses done in pain-related tissues contrasted with the genes that are simply highly expressed in those tissues, and found that the former sets of genes decoded better GWAS of pain.

## MATERIALS AND METHODS

### GENE EXPRESSION OMNIBUS DATASETS

Data was downloaded from the Gene Expression Omnibus (GEO) database deposited there up to August 2021^9,10^. We searched for the keyword “pain” in “Geo DataSets”, then restricted the study type to either “Expression profiling by array” or “Expression profiling by high throughput sequencing”. Also, we confined “Organism” to either “Mus musculus” (mouse), “Rattus norvegicus” (rat), “Sus scrofa” (pig), or “Homo sapiens” (human). We then required that the keyword “pain” appeared on the main accession page of the study. The research abstract was carefully read and if relevance and eligibility were found, the data was selected for further analysis.

### PROCESSING OF MICROARRAY DATA

The processing of microarray data was performed in R using GEOquery version 2.40.0^11^, and limma version 3.26.8.^12,13^ was used for the detection of differential gene expression. Microarray files obtained from MSigDB^14^ and Gene Expression Omnibus^9,10^ enabled the translation of microarray probe names to the study’s host specie gene symbols. Host species gene symbols were converted to humanized HUGO IDs from translation tables obtained from Ensembl’s BioMart online web tool^15,16^.

### PROCESSING OF RNA-SEQ DATA

The RNA-Seq data were downloaded from GEO using the “prefetch” tool, part of the SRA Toolkit(http://ncbi.github.io/sra-tools/). Once downloaded, the data were converted from the SRA format to FASTQ using the “fasterq-dump” program. We concatenated the FASTQ data for samples that were deposited scattered across many SRR accession IDs. Attention was paid to the sequencing style, either single- or paired-ended, and preserved at the moment of genomic mapping. The sequencing data was then mapped on the appropriate genome using STAR^17^, with the “--quantMode GeneCounts” mode enabled. Genomes were downloaded from Ensembl’s FTP site^18^, version 103; http://ftp.ensembl.org/pub/release-103/fasta/. There, the mouse genome assembly version was GRCm39, Rnor6 for the rat, and GRCh38 for humans. There were no reference genome used for Sus scrofa (pig) as the only pig dataset was from microarray technology. Differential expression of genes was detected with the help of the “DESeq2” R package^19^. Sex was used as a co-variable for studies containing animals or humans of both sexes. The age of participants was also used as a co-variable for human studies when provided. Finally, Ensembl gene accession IDs were converted to humanized HUGO IDs from translation tables obtained from Ensembl’s BioMart online web tool^4,5^.

### GENE SYMBOLS

Gene symbols were updated to current ones using translation tables kindly provided by the HUGO Gene Nomenclature Committee (HGNC)^20^. For microarray data, we first checked if a gene symbol was deprecated, and updated if needed. For RNA-Seq data, because we used a recent genes definition file during reads mapping on the various genomes, we first checked if a gene symbol was current (as opposed to deprecated) and updated it only if not found. Unfortunately, some gene symbols could be found simultaneously in both current and deprecated lists, like *VARS2*, and thus some ambiguity might have been introduced.

### PROCESSING OF PATHWAYS

The results of the differential expression of genes analysis were given for input to ‘fgsea’, where genes were sorted by decreasing test statistic; from most up-regulated to most down-regulated (https://doi.org/10.1101/060012;https://github.com/ctlab/fgsea). Pathways were from Gene Ontology,^21^ conveniently provided in the Bader lab and accessed on September 1st, 2021 (http://download.baderlab.org/EM_Genesets/September_01_2021/Human/symbol/). The human version of pathways was used for pathway analysis, as all DEGs were in the humanized symbols form.

### PAIN GENES COLLECTION

We compiled a list of genes, named “Pain Genes Collection”, found to be implicated in pain processes. These genes are coming from various sources including Pain Research Forum (https://www.painresearchforum.org/resources/pain-gene-resource), Gene Ontology for the “pain” keyword (http://amigo.geneontology.org/amigo/search/bioentity?q=pain), and Pain Genes Database^22^. The description of the sources and the complete known pain gene list could be found in Supplementary Material (Supplementary Table S1).

### IDENTIFICATION OF TRANSCRIPTIONAL PAIN BIOMARKERS CANDIDATES USING MACHINE LEARNING

Several tools from machine learning approaches were combined to unveil RNA markers capable of identifying a sample’s pain state (Supplementary Figure S1). We utilized the machine learning approach built into a tool named Evaluation of Predictive Capability for ranking biomarker candidates (EPCY), which performed leave-one-out cross-validations (https://doi.org/10.7490/f1000research.1117272.1). We used the Matthews correlation coefficient (MCC) performance metrics from EPCY since we have an acute class imbalance for pain classes^23^. EPCY reported the MCC for each gene from a contingency table filled by predicting the class of the left-out samples, such that the MCC is for unseen examples throughout, in a way that the learning algorithm is applied once per instance, with all other instances serving as a training set and the selected instance serving as a single-item test set. For each of the four pain classes, namely naïve, sham inflammatory, and neuropathic pains, we first asked EPCY to distinguish a pain class from all others (e.g. naïve versus not naïve). This yielded four lists of genes, one for each pain class, in which genes were sorted by decreasing MCC (from most correlated to least) (Supplementary Figure S1A). Using the ‘caret’ package in the R statistical software (https://CRAN.R-project.org/package=caret), we performed logistic regressions between a pain class and gene expression for the top two, three, …, up to the top 50 genes to force overfitting (Supplementary Figure S1B). This way, we could track Cohen’s kappa performance metric on 10-fold 100-repeat cross-validations. In caret, kappa is reported instead of the MCC for test set performances by default, even though kappa is appropriate for unbalanced class counts, it was found to have undesired behavior in special circumstances (not applicable here)^24^. As the number of genes considered in the regressions increased, the prediction performance increased, hit a maximum, then started to decrease (evidence of over-fitting). The number of genes T at the maximum performance was then used in a final round of LASSO with these top T genes, using again the kappa performance metric on a 10-fold 100-repeat cross-validations (Supplementary Figure S1C). LASSO was used as it would detect co-varying genes, removing any redundant genes from the final list of predictors (LASSO-assigned weight of zero), thus providing for a most parsimonious model(https://www.jstor.org/stable/2346178). We then applied the same approach for distinguishing pain states in a pair-wise manner.

### PARTITIONED HERITABILITY OF HUMAN CHRONIC PAIN CONDITIONS IN PAIN-RELEVANT TISSUES

The human genome was partitioned into tissue-enriched (TE) expressed genes, highly expressed (HE) genes, or DEGs following a pain assay. For TE expressed genes we used the human gene expression database published by Benita et al.,^25^ downloaded from http://xavierlab2.mgh.harvard.edu/EnrichmentProfiler/Datasets.html, which contains pain-relevant tissues such as dorsal root ganglia (DRG) and spinal cord (SC). Here, the genes were scored for their tissue specificity based on their expression level. The top 1,000 scored genes of each tissue were selected as human TE genes for that tissue. For rodents, we used HE genes in SC and DRG samples from TPSDB as a database equivalent to the human gene expression database doesn’t exist for pain related tissues. We selected top 1,000 genes with the highest average transcript per million (TPM) value in that tissue.

For DEGs, we considered DRG or SC tissues regulation in the inflammatory and neuropathic pain assays. For each pain-tissue combination, we ranked all genes by their average absolute values of differential expression test statistics across contrasts. For each tissue, we extracted the top 1,000 genes and used them for partitioning heritability using LDSC^26,27^. Partitioned heritability analyzes were performed for two human chronic pain phenotypes from a study by Khoury^28^. First, for single site chronic pain (SSCP) from a UK Biobank GWAS comprised of 175,769 individuals (93,964 cases and 81,805 controls). Cases were defined as participants who report only one chronic pain site and controls were defined as participants who answered: “None of the above” to data field 6159 in UK Biobank cohort. Second, for multisite chronic pain (MSCP) from a UK Biobank GWAS comprised of 164,778 individuals (82,812 cases and 81,966 controls), where cases were defined as participants who report at least two chronic pain sites while controls had the same definition as in CLPC group.

### TPSDB WEBSITE

For the building of the website, a web app framework called Django was used (https://www.djangoproject.com/). The framework takes the HTML webpages and uses python to process and present the web pages. This is done using views, which are able to process the HTML and python to dynamically return web pages.

## RESULTS

### TPSDB WEBSITE

For easier use of the TPSDB, a website design enables users to search the database. On the homepage, after selecting the search box of interest (Gene, Pathway, or SNP), the website takes a post form with the requested item (Supplementary Figure S2). Then, to request results from the database, the user must type the name of their gene, pathway, or SNP. Users can also filter the results based on a variety of criteria, such as the host’s sex, species, tissue, and the type of pain response it is experiencing, among others (See Supplementary Figures S3). For the tissue filter, the user has to use the Medical Subject Headings (MeSH) code from the National Library of Medicine (https://www.nlm.nih.gov/mesh/2019/download/2019New_Mesh_Tree_Hierarchy.txt) (See Supplementary Figures S4).

Once the user requests a search, the results will be displayed as a table. This table contains all the results related to their request and can be further searched using a search box located in the upper right corner of the table (Supplementary Figure S3B). Clicking on each column’s name in the "Column Description" part at the top of the result box will reveal the description of that column. The Excel file of the results for offline viewing is downloadable by clicking "Download Your Results" (Supplementary Figure S5).

### VARIETY OF GENE EXPRESSION COMPARISONS IN TPSDB

At the creation of TPSDB, 338 DEGs’ contrasts have been done (Supplementary Table S2). A number of pair-wise contrasts have been made in each available transcriptomics dataset ranging from 1 to around 50 (Figure 1A). In terms of gene expression measurement method, over half of the contrasts in the database are from high throughput sequencing (HTS) and the rest were from microarray (ARRAY) (Figure 1B) and the HTS proportion increases with time (Figure 1C). Most of the contrasts have been done in datasets from rodents (mouse and rat) and the rest are from human and pig datasets respectively (Figure 1D). The most frequently assessed tissues were of the peripheral and central nervous systems origin (e.g. DRG, brain)(Figure 1E). The considered pair-wise contrasts were mostly concerning to comparison of pain state versus control (Figure 1F), followed by gene expression changes over time and then by sex-related differences. Historically, the majority of the studies have been done in males only, but mixed male/female and female-only studies are also growing overall (Figure 1G) (Supplementary figure S6). The samples that we used cover a variety of pain types that mostly could be combined under the umbrella of neuropathic or inflammatory pain (Figure 1H,I). Named pain conditions are either related to a disease (e.g. arthritis) or are related to animal pain assays (e.g. spared nerve injury, sciatic nerve ligation; Figure 1I). Considering tissue and specie-wise analyses (Supplementary figure S7), blood-derived studies were mostly comprised of humans, while studies in the nervous system were done about 2/3 in mice and 1/3 in rats.

**Figure 1.**
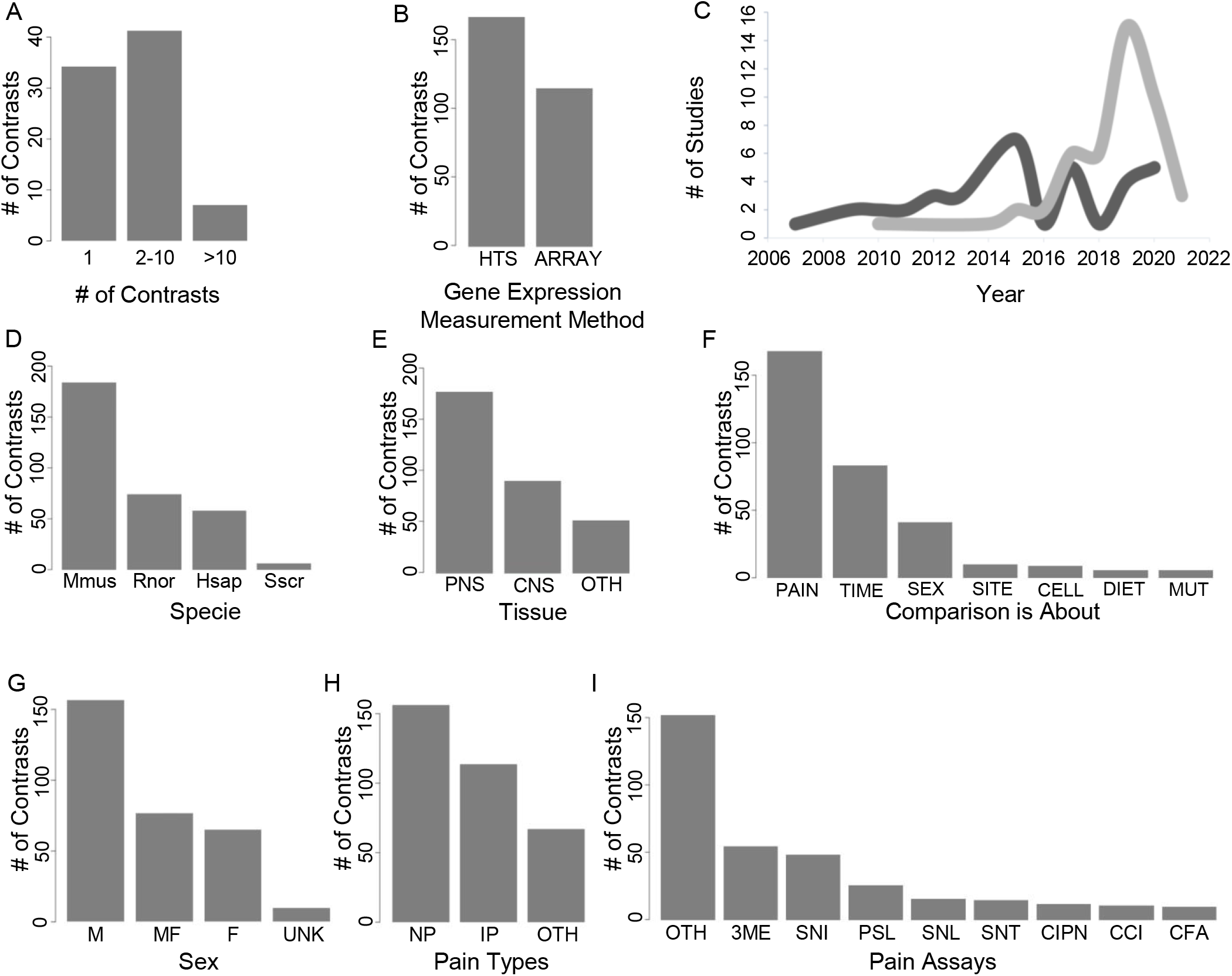
TPSDB covers a wide variety of contrasts of gene expression. (A) Number of studies per number of contrasts. (B) Number of contrasts as a function of gene expression measurement method: high-throughput sequencing, HTS; cDNA microarray, ARRAY. (C) Usage of each gene expression measurement methods over time. The y-axis shows the number of datasets in that year, while the x-axis represents the year. Dark gray represents datasets that have used microarray method. Light gray represents datasets that have used high throughput RNA sequencing method. (E) Number of contrasts as a function of the anatomical location of the extracted tissue: peripheral nervous system, PNS; central nervous system, CNS; and others. (F) Number of contrasts per type of comparison: pain assays or pain condition, PAIN; time points in each pain assay, TIME; sexes, SEX; cell types, CELL; pain sites, SITE; diet, DIET; mutated genome, MUT. (G) Number of contrasts based on the sex of specie used in the study: males-only, M; females-only, F; males and females combined, MF; unknown, UNK. (H) Number of contrasts based on the pain type: neuropathic pain, NP; inflammatory pain, IP; and other pein types, OTH. (I) Number of contrasts per pain induction method: other than mentioned, OTH; third molar extraction, 3ME; spared nerve injury, SNI; partial sciatic nerve ligation, PSL; spinal nerve ligation, SNL; sciatic nerve transection, SNT; chemotherapy-induced neuropathic pain, CIPN; chronic constriction injury, CCI; complete Freund’s adjuvant, CFA.

### GENES WITH THE HIGHEST EVIDENCE FOR DIFFERENTIAL EXPRESSION FROM THE TPSDB

Using the results of DEGs analyses across all the contrasts, we ranked genes by the frequency of being differentially expressed in TPDB (Supplementary Table S3). The top 10 genes with the highest ranking of being differentially expressed in the majority of contrasts, significantly up or down-regulated in a tissue-specific manner are presented in Figure 2A-D (left panel). Importantly, for the genes with the highest differential expression, the proportion of the contrasts where their differential expression can be detected at a significant level is very high and ranges from 34% in blood to 65 % in the sciatic nerve (SN), suggesting the shared mechanisms of gene expression regulation in different species and pain states^29^.

**Figure 2.**
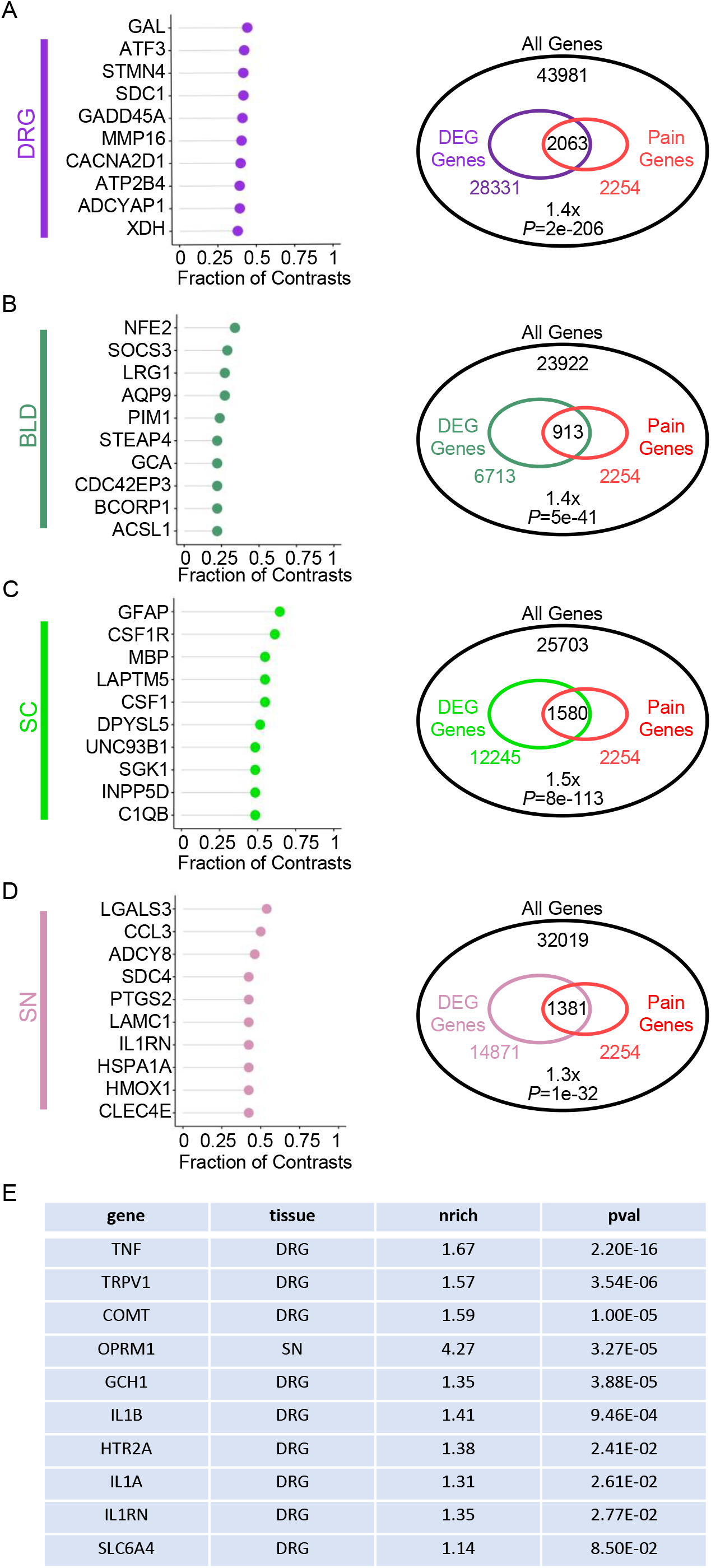
Gene-level statistics from the TPSDB. (A-D) Top ten genes most significantly differentially expressed (left). Plots track the proportion of contrasts per tissue where each gene was significantly up- or down-regulated at the FDR 10% level. Overlap of all tissue-specific significantly differentially expressed genes with the 2254 known pain genes in “Pain Gene Collection” (right). Per each tissue, Venn diagrams show the total number of detected genes, the total number of up- or down-regulated genes at the FDR 10% level (DEG), overlap with known pain genes, the enrichment fold at the Venn intersection along with the associated P-value for enrichment. Tissues are: (A) dorsal root ganglia (DRG); (B) whole blood (BLD); (C) spinal cord (SC); and (D) sciatic nerve (SN). (E) The enrichment for of “popular pain genes” in TPSDB with the tissue that they were enriched in.

We next tested the overlap of significant DEGs (FDR 10%) with the 2254 genes already implicated in pain (“Pain Gene Collection”). Overall, the enrichment was high in all the tissues with the highest enrichment, overlap, and statistical significance in DRGs and the lowest overlap and number of DEGs in blood, although it had the second greatest number of samples in the database (Figure 2A-D, right panel). Overall, and predictably, the unbiased transcriptome-wide assessment of pain-related DEGs identified almost an order of magnitude more pain-related genes than previously known.

To characterize the differential expression of the genes that were reported as historically the most “popular pain genes” before^30^, we tested their enrichment score in our dataset in different tissues. We observed significant enrichment for all of them (Figure 2E). The fold change distribution of *TRPV1* differential expression is shown as an example (Supplementary Figure S8). With exception of the Mu opioid receptor *OPRM1*, all of them were enriched in DRGs (Figure 2E; Supplementary Table S4). This can be a reflection of our sample size, which was the largest for DRGs, and the overall focus of pain research on DRG. Mu opioid receptor *OPRM1* demonstrated the highest enrichment in the SN. This can be suggestive of the particular importance of *OPRM1* expression in SN in the context of pain assays. Overall, although the unbiased transcriptome-wide assessment of pain-related DEGs validates previous related knowledge, there are many more pain-related genes of seem to be a much higher magnitude of effect size than those traditionally assessed.

### PATHWAYS WITH THE HIGHEST EVIDENCE FOR DIFFERENTIAL EXPRESSION FROM THE TPSDB

Using the results of DEGs analyses across the contrasts, we then used Gene ontology (GO) enrichment analysis to build and rank all biological pathways by the frequency of being differentially expressed in TPDB in a tissue-dependent manner (Supplementary Table S5). We analyzed biological processes (GO:BP), cellular component (GO:CC), and molecular function (GO:MF). The top 10 pathways of the BP that were of highest enrichment in DRGs (Figure 3A), blood (Figure 3B), SC (Figure 3C), and SN (Figure 3D) are shown. Regardless of tissue, cytokine related pathways were among the top 10 enriched biological processes. Innate immune response was also highly regulated in both blood and SC tissues. The enrichment analysis results for the top 10 cellular components are suggestive of the importance of mitochondrial membrane, being the only cellular component enriched in all the tissues (Supplementary Figure S9). Ion channel complex and its activity were the cellular components and molecular functions, that were specifically enriched in central and peripheral nervous system tissues including DRG, SC, and SN (Supplementary Figure S9 and S10)(Supplementary table S5).

**Figure 3.**
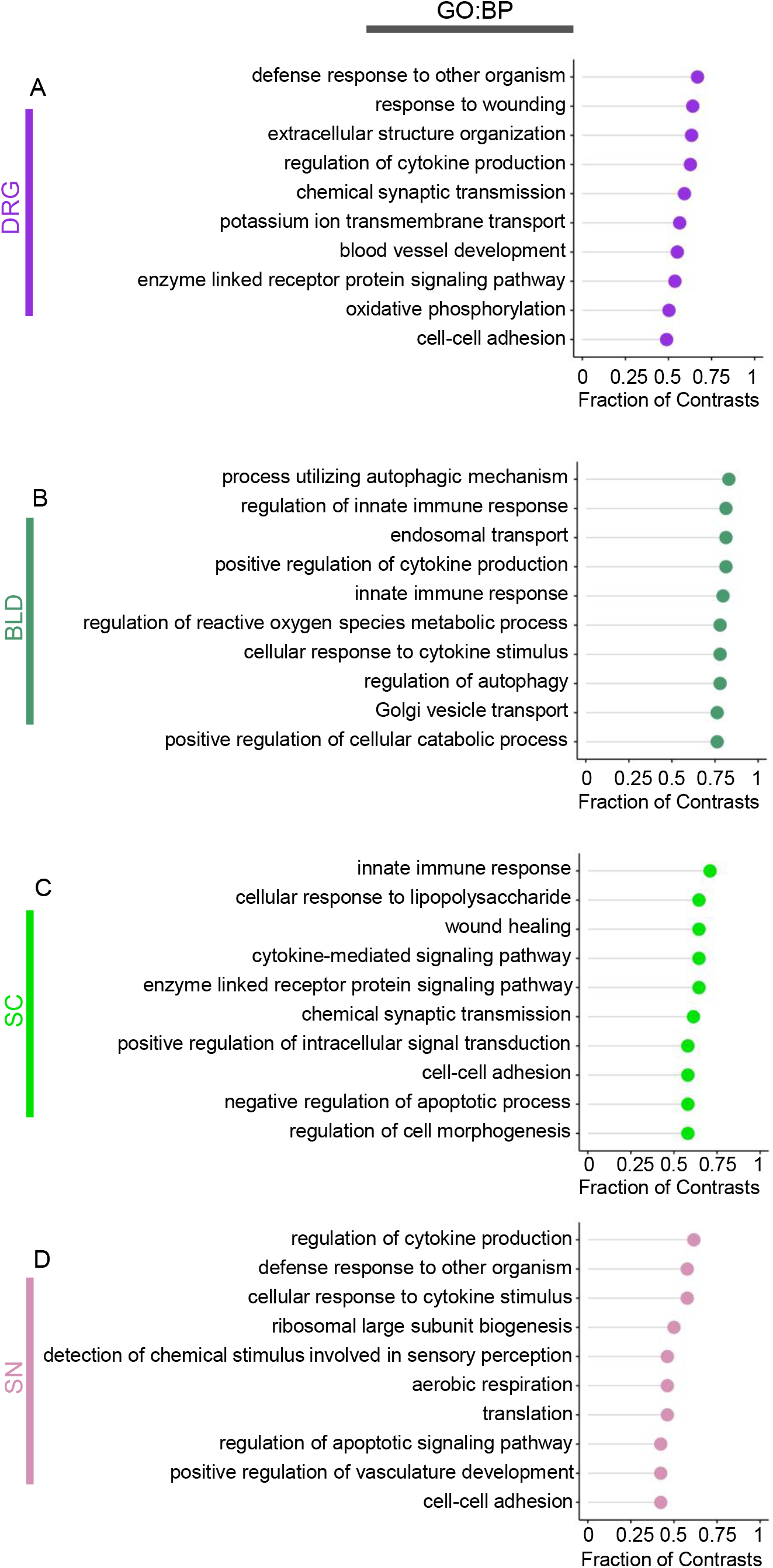
Pathway-level statistics from the TPSDB database. Top ten biological processes (GO:BP) most significantly enriched per tissue. Plots track the proportion of contrasts per tissue where each pathway was most represented in enrichment analysis. (A) dorsal root ganglia (DRG); (B) whole blood (BLD); (C) spinal cord (SC); and (D) sciatic nerve (SN).

### PAIN STATE RNA CANDIDATE BIOMARKER FROM TPSDB

We next wanted to take advantage of a large and diverse transcriptomics database to find RNA markers to differentiate pain states using a machine learning approach. We tested if the four pain states, namely naïve, sham, inflammatory, and neuropathic pains, could be distinguishable by the expression levels of a handful number of genes (Supplementary Figure S11). For this, we analyzed samples from rodents (mouse and rat) in DRG; a combination that offered the greatest number of samples (a total of 141; 37 naïve, 41 sham: 8 IP sham and 33 NP sham, 18 IP, and 45 NP).

At the first stage, we asked for each pain state to distinguish “being a member of a specific pain state” from “not being a member of a specific pain state” using a combination of tools from machine learning. Using Landis and Koch’s strength of agreement for various kappa ranges we could interpret the classification performance based on gene expression TPMs (Table 1)^31^. Based on the established criteria, kappa values in the range 0-0.20 are considered as slight, 0.21-0.40 as fair, 0.41-0.60 as moderate, 0.61-0.80 as substantial, and 0.81-1 as almost perfect. All four pain states could be predicted in a fair fashion or better, with the worst average kappa of 0.36 for inflammatory pain with 18 genes at maximum kappa. The sham state, consisting of both neuropathic and inflammatory pain shams, could be moderately well predicted with average kappa of 0.58 using 18 genes at maximum kappa. The higher number of genes at maximum kappa needed for prediction and the lower kappa values indicated that the sham and inflammatory pain states are harder to accurately predict. Substantial prediction performances were observed for naïve and neuropathic pain with average kappa of 0.71: we needed only two predictor genes for the naive state and 13 predictor genes for NP. This might be due to the fact that sham interventions elicit inflammation in a manner similar to inflammatory pain assays.

**Table 1.**
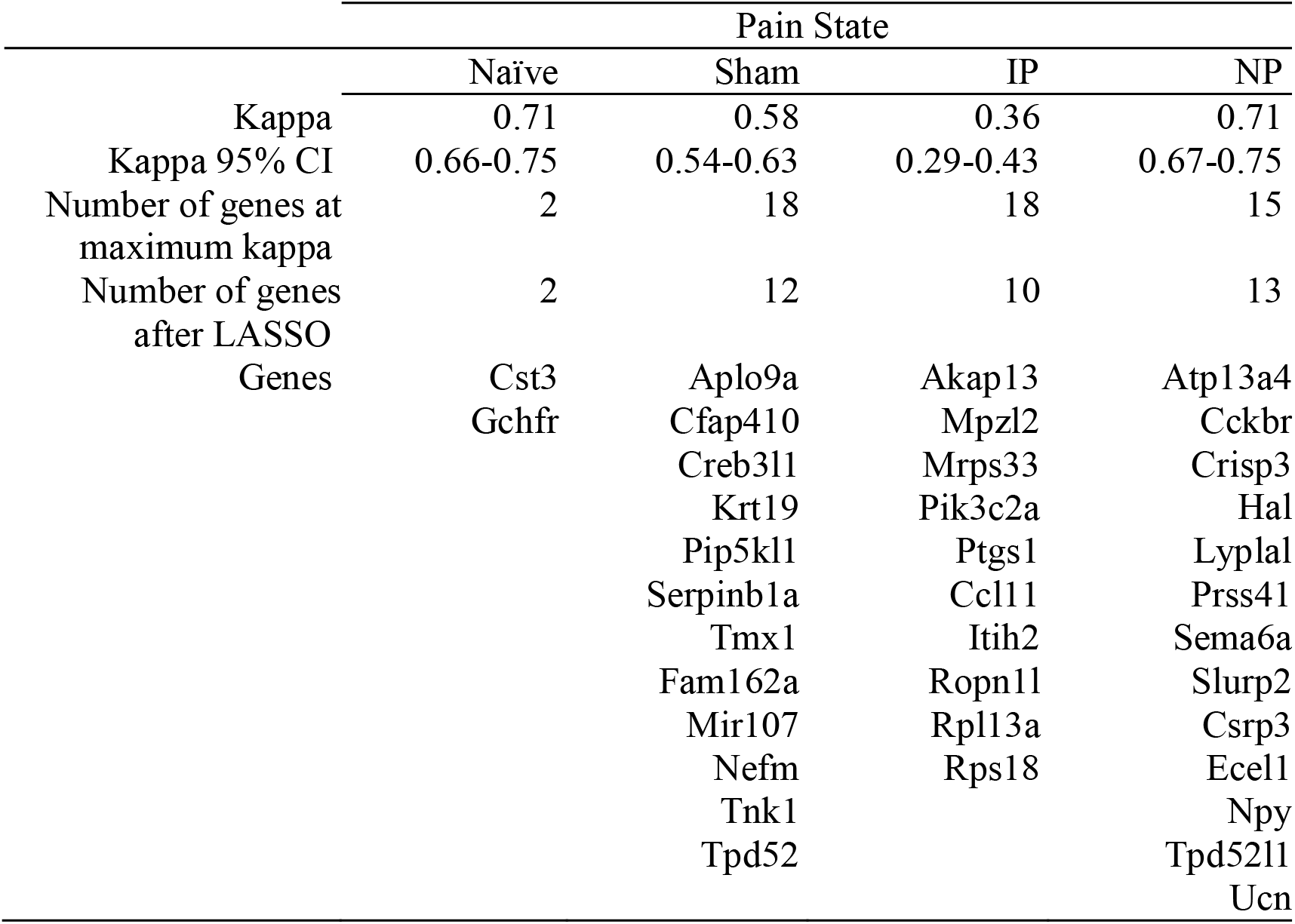
Identification of biomarker candidates to differentiate four pain states between each other: naïve, sham, inflammatory pain(IP), and neuropathic pain (NP) in rodent dorsal root ganglia.

Then we applied the same approach for distinguishing pain states from their controls and the neuropathic and inflammatory pain states themselves, in a pair-wise manner, to pinpoint precisely which sample groups are hindering the accurate detection of state-specific markers. We separated sham of neuropathic pain (NP Sham) and sham of inflammatory pain (IP Sham) in this analysis and then we found the best predictor RNA for detection of “IP from IP sham”, “NP from NP sham” and “NP from IP”. Restricting the samples to only two pain states has led to a significant increase in Kappa and prediction ability, and as expected, a smaller number of genes for having the maximum Kappa number. Table 2 presents the results obtained from the pair-wise analysis for pain state-specific RNA marker. As can be seen, NP and IP are perfectly distinguishable using 6 genes. IP and IP sham are also perfectly distinguishable with only 2 genes. Although NP was distinguishable from its sham control in a substantial manner, it had the lowest kappa value of 0.62 among others meaning that NP and NP sham samples were harder to distinguish.

**Table 2.**
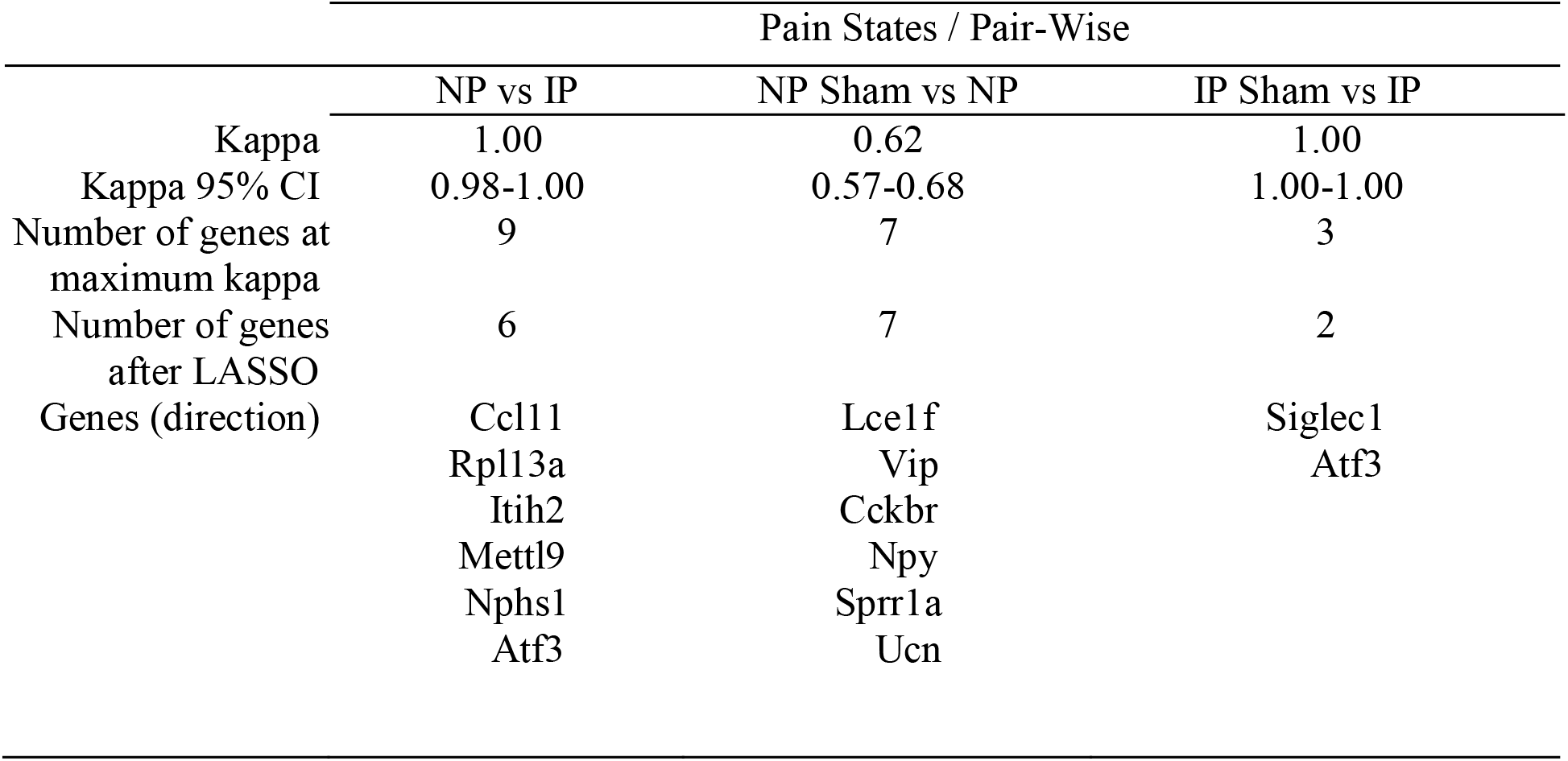
Identification of biomarker candidates to differentiate pain states in a pairwise comparison in rodent dorsal root ganglia: naïve, sham, inflammatory pain(IP), and neuropathic pain (NP).

### PARTITIONED HERITABILITY OF HUMAN CHRONIC PAIN CONDITIONS IN PAIN-RELEVANT TISSUES

We next employed our large transcriptomics dataset for functional analysis of GWAS results. We tested how partitioned heritability enrichment at loci of tissue-specific genes at their base expression level in a pain-relevant tissue would compare to that at loci of genes differentially expressed following a pain assay in the same tissue (Figure 4). To do so, we first extracted genes from human and rodent DRG and SC at their baline expression, namely, tissue-enriched expressed genes in humans (TE) and genes highly expressed (HE) in rodents. We then performed partitioned heritability in two human GWASes of chronic pain, single-site chronic pain and multisite chronic pain (Figure 4). There, we did not find evidence for enriched heritability at genes expressed in either DRG or SC. This was despite the many studies that demonstrated the contributions of SC and DRG to chronic pain, and the non-negligible narrow-sense heritability estimates for single-site and multisite chronic pain conditions (6.9% for the former, 19.1% for the latter)^32^. We next extracted genes most differentially expressed in inflammatory and neuropathic pain assays in DRG and SC. We found that loci at genes differentially expressed in DRG after neuropathic pain assays yielded a significant heritability enrichment (FDR=20%) for multisite chronic pain (Figure 4B). We did not find significant heritability associated with genes differentially expressed in SC after neuropathic pain assays, nor in DRGs or SC after inflammatory pain assays. Our results suggest that human multisite pain is driven mostly not by just DRG-specific genes, but by genes differentially expressed in DRG in relevance to neuropathic pain. Finally, we did not find evidence for enriched heritability at genes expressed in either DRG or SC for single-site chronic pain (Figure 4A), which is probably a reflection of the lower heritability of single-site chronic pain.

**Figure 4.**
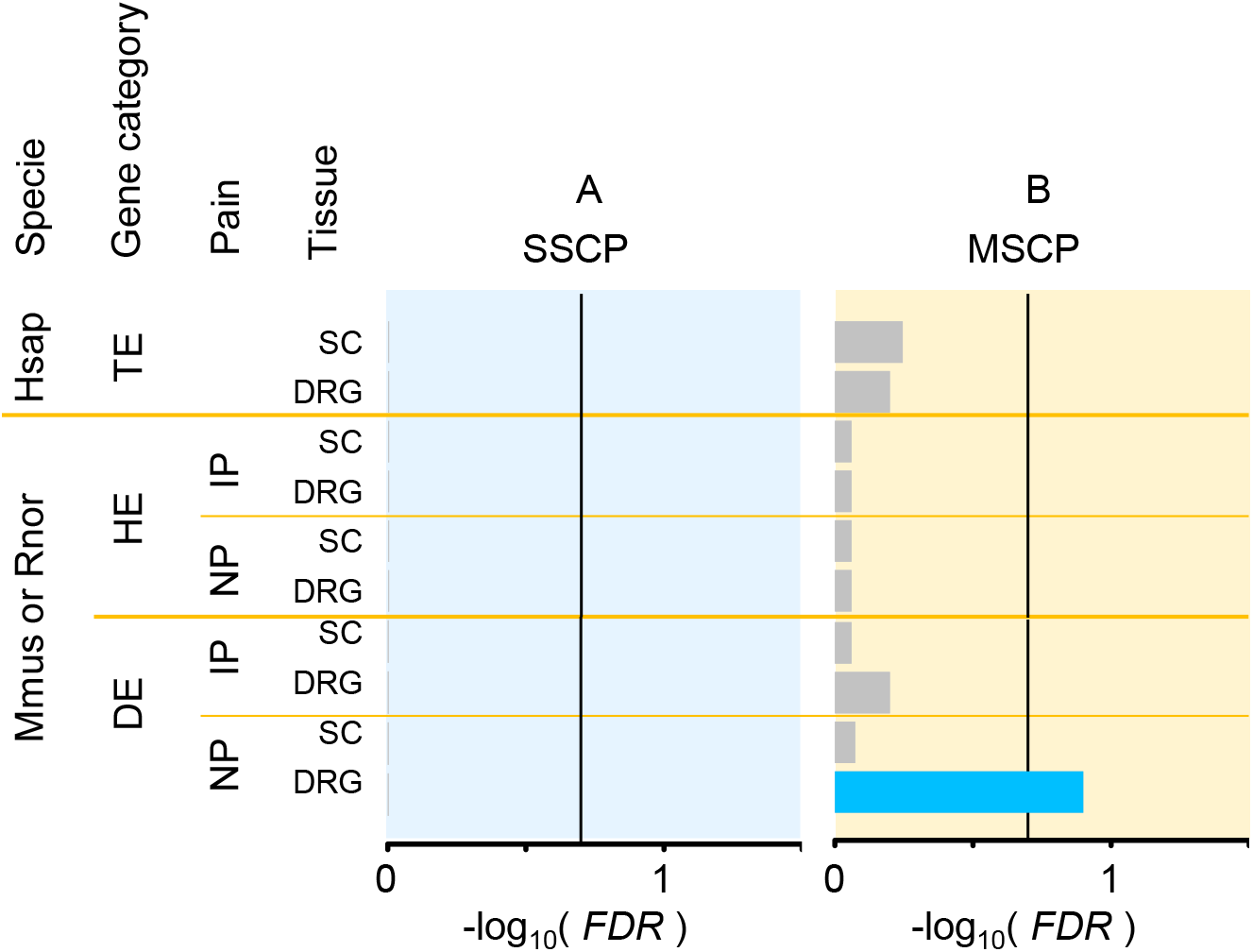
Partitioned heritability of GWAS results of multisite and single site human chronic pain conditions in spinal cord (SC); and dorsal root ganglia (DRG). Gene expression levels were considered in humans (Hsap), or in mice (Mmus) or rats (Rnor). The human genome was partitioned into loci of tissue-enriched expressed genes in humans (TE), genes highly expressed (HE) in a rodent tissue, or genes differentially expressed (DE) following a pain assay. Pain assay classes were: inflammatory pain (IP) or neuropathic pain (NP). Barplots show FDR-corrected P-values for enrichment of heritability at loci of genes of interest. Bars filled in grey when not significant or blue otherwise. Vertical black bar indicates statistical significance at the FDR 20% level. (A) Partitioned heritability in single-site chronic pain conditions (SSCP). (B) Partitioned heritability in multisite chronic pain conditions (MSCP).

## DISCUSSION

The TPSDB is introduced as a resource for accessing the result of DEG analysis from the multiple publicly available transcriptomics datasets related to pain. Here we describe the results of the 338 contrasts for full transcriptomics datasets for the first version of the database (Supplementary Table S2). It is made to let researchers to benefit from the wealth of ample amount of publicly available data, at the same time, and in a convenient way. We aimed to preserve the variety of the transcriptomes sources as much as possible by having different pain phenotypes, organisms, tissues, and time points. The approach here was towards non-biased analysis of all pain related Microarray and RNA-seq studies available on GEO, making a comprehensive resource for checking on a hypothesis for specific genes, SNPs, and/or pathways or downloading the raw data and performing a hypothesis-free analysis.

Numerous studies have compared gene expression quantification via microarray versus RNA-Seq^33,34^. In the microarray approach, RNA quantification is achieved via in vitro hybridization, between actual RNA fragments and the microarray’s probes. In RNA-Seq, RNA quantification is achieved in silico. As such, microarrays are more sensitive to genetic content at probed loci. However, RNA-seq better discriminates between highly homologous sequences and allows the discovery of new genes. Thus, both of these methods have strengths, provide useful information on gene expression on a genome-wide scale, and there are multiple publicly available datasets produced using both these technologies, therefore we utilized both data types to create TPSDB.

It is important to keep in mind that each transcriptomics dataset comes with its assay characteristics such as batch effects and sequencing depth^35^. To address these possible problems coming from the above mentioned variability, we kept each set of the differentially expressed analysis within each study and did not mix any samples of the same or similar conditions. Moreover, RNA-Seq data enables further transcriptomics analyses, in particular, differential expression of exons (e.g. DESeq^19^) and isoforms^36^. It also enables the detection of new isoforms, either from retained introns or exons not known to be transcribed. RNA-Seq (and recent microarray) also allows for the study of non-coding transcripts^37^. We plan to include these extra analyzes in future releases.

Another difficulty of doing transcriptomics studies is the fact that results will have a probably long list of DEGs sorted by their statistical significance, but the relative biological importance of each gene may be unclear, as some of these genes might be highly ranked but related to factors other than our favorable perturbation. One way to address it is to look over many experiments and datasets related to the same phenotype with slight variations and consider the number of times that each gene appears among significant DEGs.

In this study, we did a number of analyses that help us both to characterize our TPSDB and start to utilize the power of multiple transcriptomics datasets. First, we performed the counts for both genes and pathways that are regulated in most of the pain phenotypes regardless of the pain type, time point, data type, and organism. At the gene level, in DRG we found *GAL* as the most repeated DEG which codes a neuropeptide with known and important roles in both acute and chronic pain and that was already suggested as a potential therapeutic target in pain conditions^38^. As the second ranking gene in DRG, we found a transcription factor *ATF3* with a known role in axonal regeneration in both the central and peripheral nervous systems. This gene is also suggested to be necessary for transcriptional reprogramming and regeneration of DRG neurons after injury^39^. The rest of the genes significantly and repeatedly differentially expressed in DRG, included both new and well-known pain genes. *CACNA2D1, GADD45A, ADCYAP1*, and *XDH* genes were already in our “Pain Gene Collection”, yet many of the identified DEGs were not. Among those genes, *STMN4* is a good new candidate, as it is known to be involved in neuron projection development^40^. *SDC1* plays a role in immune response regulation by contributing to cytokine and chemokine gradients in choroid plexus epithelial barrier in brain^41^. So, our dataset is a rich source of new potentially important genes contributing to pain, especially among the genes repeatedly regulated in many tissues in diverse pain states.

Doing a pathway count on all the contrasts revealed that cytokines are highly regulated in pain assays and pain phenotypes in all of the tested pain-related tissues, including the nervous system, which is in line with previous findings related to the important role of cytokines in nociceptive processing^42^. Immune system related pathways and response to wounding were also among the biological processes which were repeatedly seen in the most of the contrasts, not only in blood, but also in as the top repeated pathways in both DRG and SC, further highlighting the importance of neuro-immune interaction in pain development.

We also took advantage of the high number of available full transcriptomics pain data in rodents’ DRG to suggest pain state RNA biomarker candidates. Using machine learning approaches, we searched for RNA marker candidates of different pain states, merely inflammatory pain, neuropathic pain, sham controls, and naïve controls, and how these states can be differentiated from non-pain states at the transcriptomics level. Among the rodent DRG pain markers that we found, some of them (*NPY, UCN, ATF3*, and *CCL11*) have been among the well-known pain genes but there were also other genes (*METTL9, ITIH2, SPRR1A*, and *KRT19*) that have never been reported as pain-related and they remain to be further studied. Many of these newly found pain biomarkers are in the same family of proteins with the ones of the known pain function. For instance, *SEMA6A* is in the same family as *SEMA3A* (Semaphorins family) which has been reported previously to have a suppressive role in peripheral nerve injury-induced neuropathic pain^43^. Same is true about *PRSS41* (Serine Protease family), *SLURP2* (Ly6/uPAR family), *AKAP13* (A-kinase anchor proteins family), *PIK3C2A* (phosphoinositide 3-kinase family), *SERPINB1A* (Serpin superfamily), and *PIP5KL1* (Phosphatide ylinositol-4-Phosphate 5-Kinase family

Although the pain-state DRG biomarker candidates identified here can have utility for *in vitro* studies and further our understanding of molecular pain pathophysiology at large, especially due to conserved somatosensation features between rodents and primates^44^, finding human blood pain biomarkers is of the most importance among our future objectives. Non-invasive nature of blood tests for biomarkers makes them proper for a variety of purposes, including pain diagnosis and quantification, better pain type classification, and monitoring treatment efficacy^45^. With an increase in the number of publicly available human datasets, we will be able to use a machine learning approach to find pain score and pain state RNA biomarker candidates.

Having access to a large number of DEGs in the context of pain provided us with the opportunity of using this set for post GWAS functional analysis, as it remains a challenge even with development of eQTL analysis techniques^46^. Partitioning of heritability at loci of genes expressed in specific cell types and tissues has already proved useful to decode the genetics underpinnings of many human conditions, like schizophrenia or auto-immune diseases^44,47,48^. Furthermore, it has been proposed that disease-associated DNA variations from GWAS studies are more likely to be found in highly regulated DEGs rather than in the genes that are selected based on their tissue-specific expression^49^. Here, we posited and tested that, similarly, genes differentially expressed in a rodent pain assay rather than just tissue-specific genes could be a better tool to decode GWAS results, thus producing an extra level of integration. In that regard, TPSDB would be complementary to our previous database which was gathering pain-related variants at the genome level^50^.

Future updates of TPSDB will provide the users with the opportunity of having access to more recent datasets and so the higher number of DEGs in the context of pain that can be used for purposes similar to partitioned heritability analysis. Since tissues are often comprised of dozens or hundreds of different cell types, transcriptome analysis at the single-cell level enabled further insights, for example, mapping genomic loci associated with human chronic pain onto primate sensory neuron (Kupari et al. 2021). Adding single-cell datasets to our database is another important future objective of database development that will let researchers to study pain transciptomics at the cell population level.

In conclusion, the TPSDB database offered here represents a comprehensive collection of pain-related transcriptome responses in various conditions and different tissues and species. TPSDB is a convenient resource for fast and easy testing on a specific hypothesis but also allows hypothesis-free research in the entire database. It will provide us with further opportunity to expand our knowledge of the molecular pathophysiology of pain by having access to all possible differential expression contrasts including those that might fall beyond the purview of each research project yet of importance for the study of pain, such as sex and time dependent transcriptional differences.

## Supporting information

supplementary tables description

## ACKNOWLEDGMENT

The current study was conducted under UK Biobank application 20802. We thank Michael Lu for prototyping the transcriptomics pain signatures database website. We would like to thank Dr. Eric Audemard for all the help with EPCY.

---

**Figure S1.**
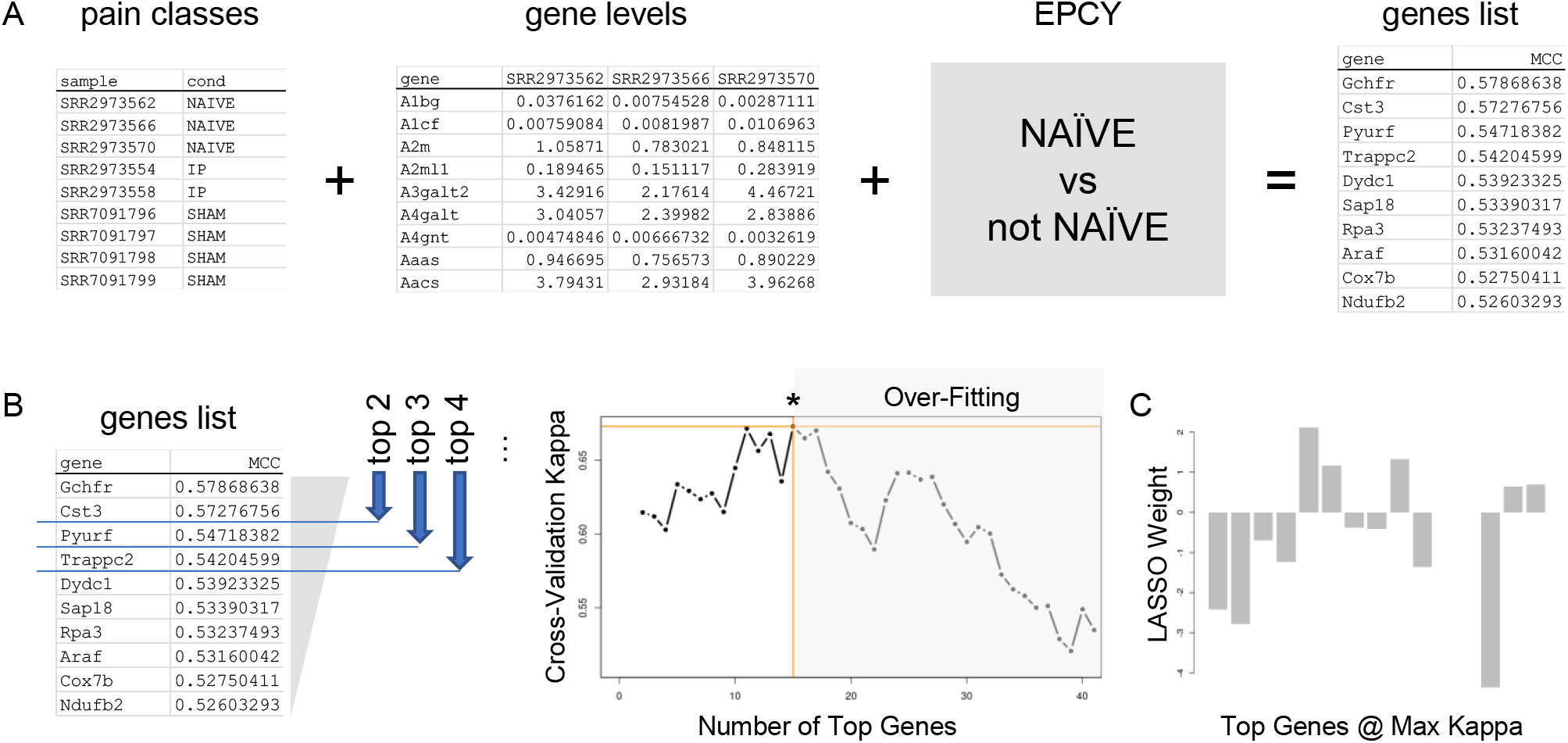
Machine learning approach for pain state marker discovery. (A) The Evaluation of Predictive Capability for marker candidates ranking (EPCY) framework. The computer program EPCY evaluated each gene’s fitness for the classification task of distinguishing a pain state (naïve for example) from all others (sham, inflammatory and neuropathic pain) based on gene expression levels in samples of different pain states. The gene prediction fitness was assessed using Matthews’ correlation coefficient (MCC). (B) Cross-validation performance measure (Cohen’s Kappa) on the test set was tracked as a function of logistic regression classification using the top two genes, the top three genes, etc. The curve displays an inverse U shape, where the maximum Kappa value is reached (indicated by ‘*’). Signs of training set over-fitting are detected by a drop in predicted performance on the test set (shaded grey area). (C) The top genes at the maximum Kappa value were submitted to a final round of cross-validation via LASSO, which assigned weights of zero to genes whose expression would co-vary with others in the top list (redundancy removal).

**Figure S2.**
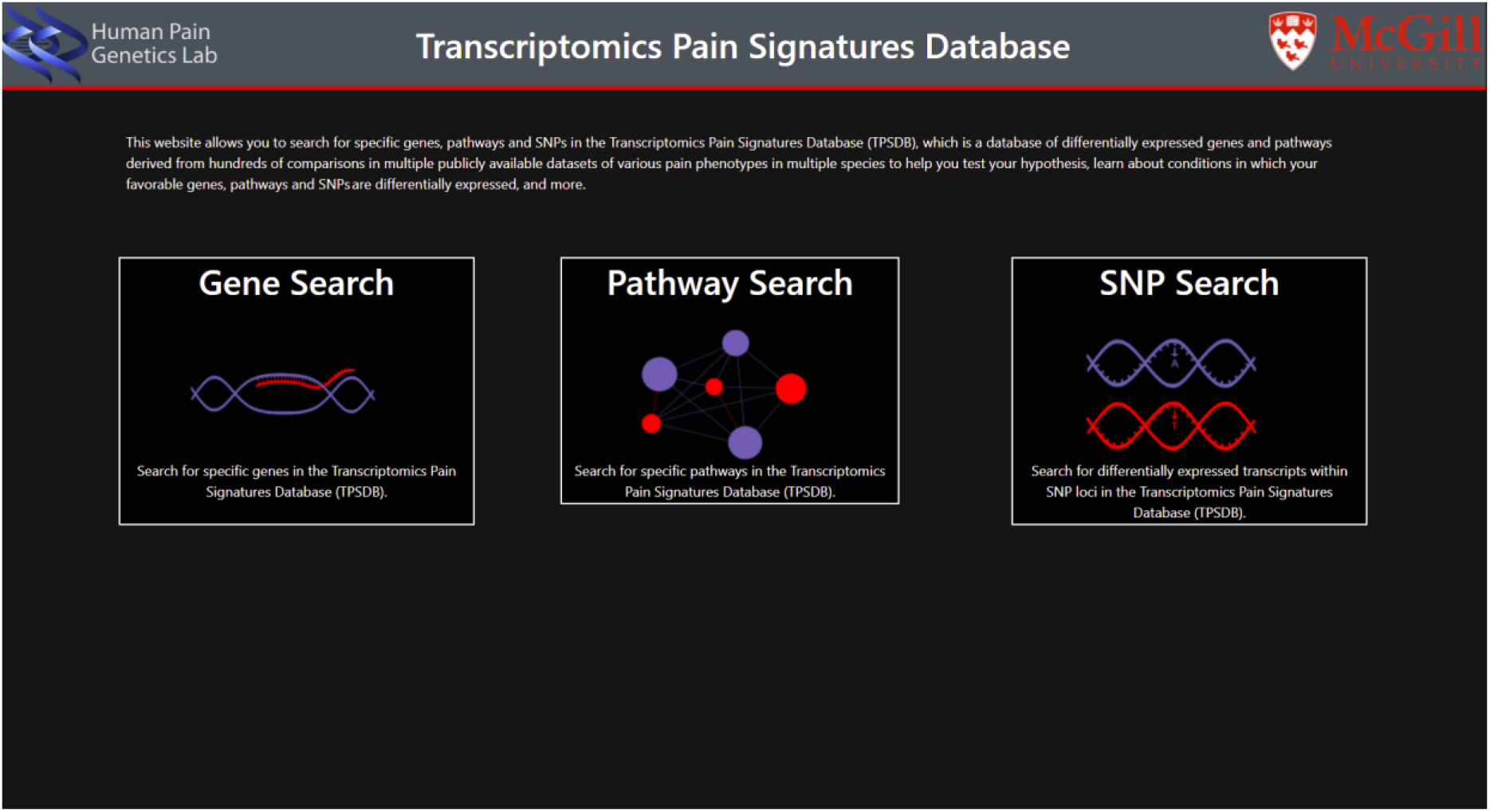
Homepage of the TPSDB website. Search box selection: depending on what is considered to be searched in the next step, the user can select gene, pathway, or SNP search box to be directed to that specific search page.

**Figure S3.**
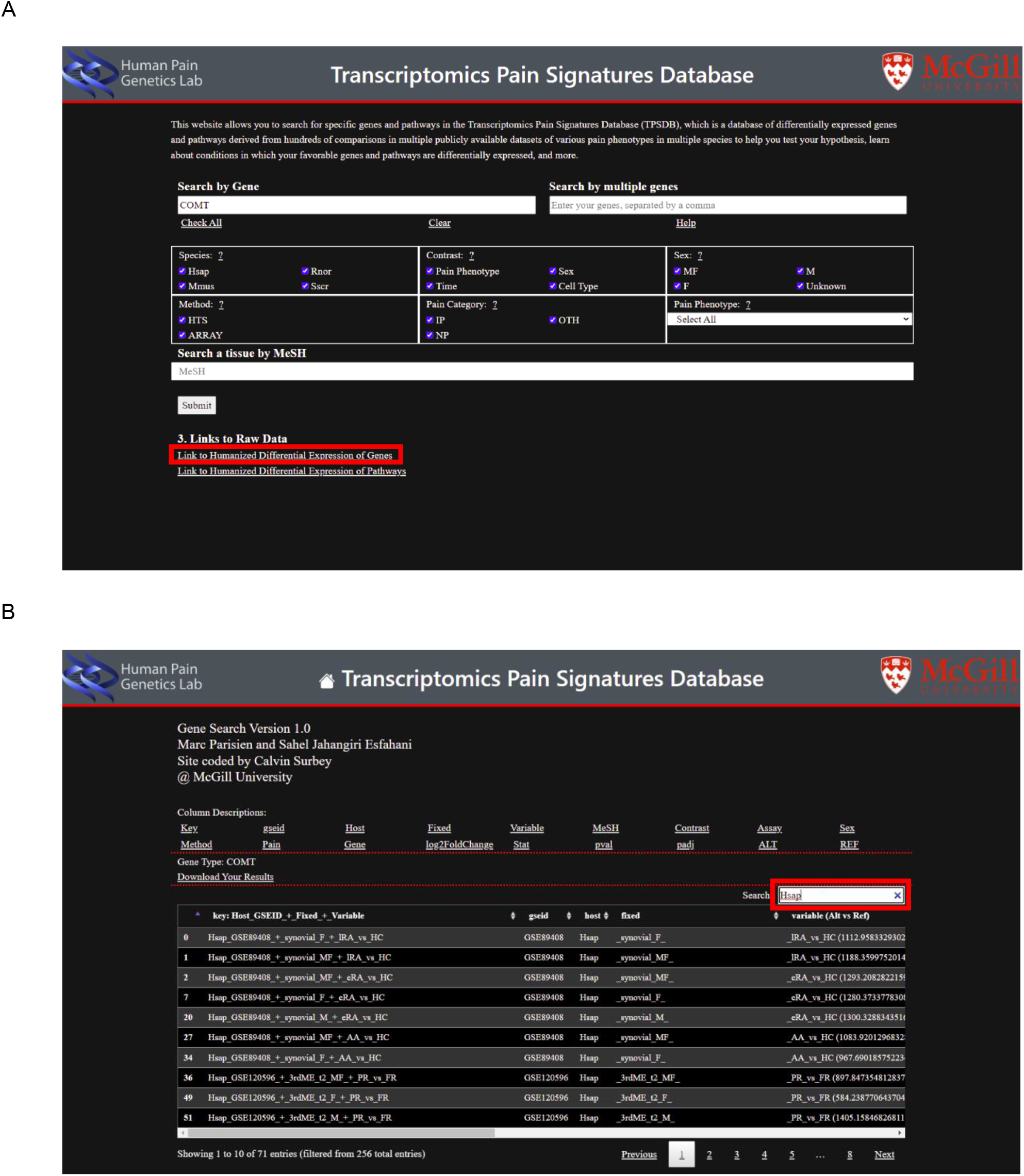
TPSDB website. (A) Gene search page. As an example, a search for the COMT gene in all sexes, species, and pain states is shown. All the results of DEG analysis available in the database are downloadable by clicking on “Link to Humanized Differential Expression of Genes” (red box). (B) The output for the differential gene expression. A search result for the COMT gene is presented as an example. Search box selection allows further refinement for the output, for example, “Hsap” (red box) is used as a search criterion which in turn will filter the results for Homo Sapiens-only contrasts. It is important to scroll to the right to see all columns of the results page as there will be 18 columns overall.

**Figure S4.**
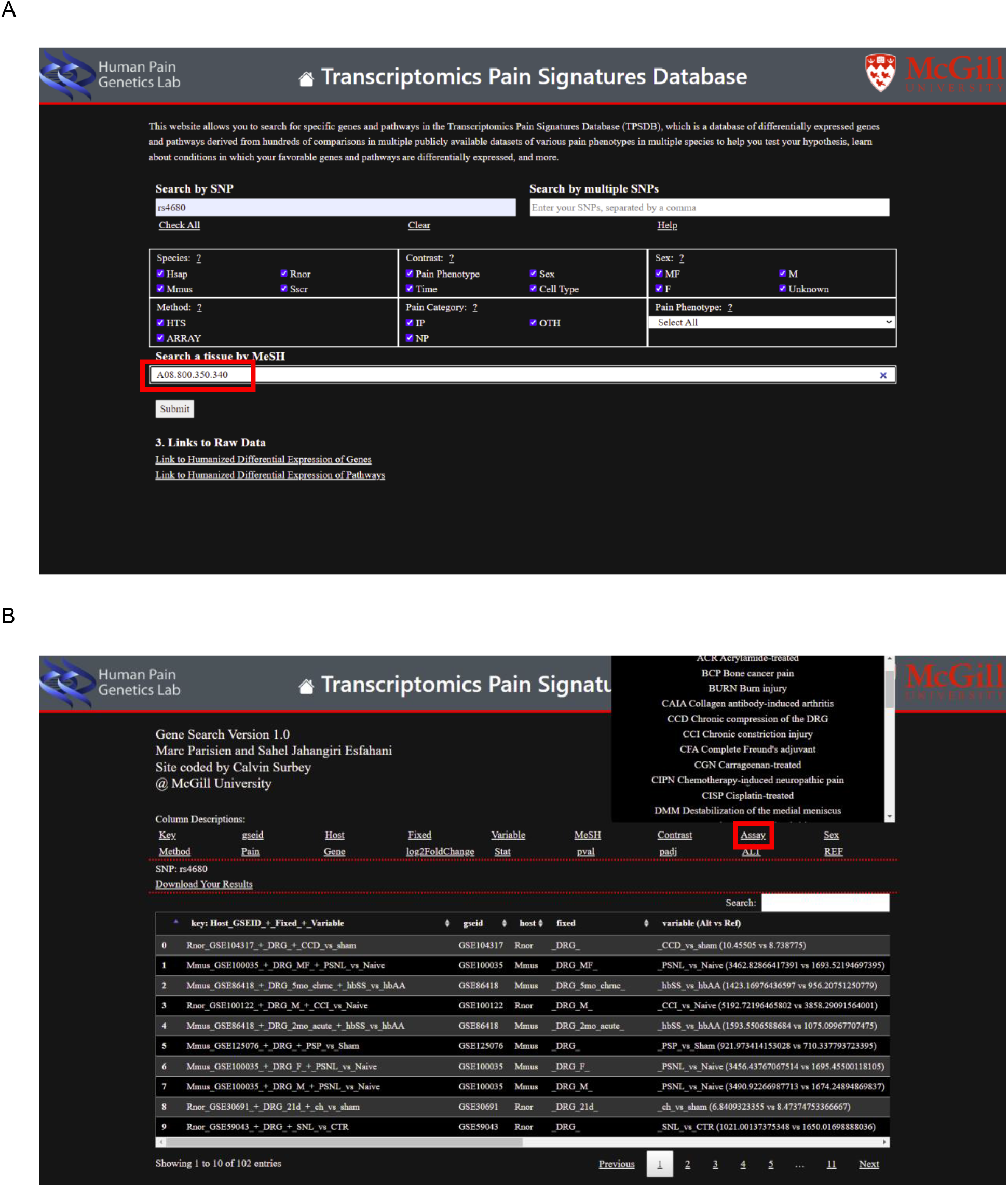
TPSDB website. (A) SNP search page. As an example, a search for SNP rs4680 within COMT gene locus in all sexes, species, pain states, and in DRG tissue (“A08.800.350.340” in the red box is the MeSH number for DRG) is shown. (B) The output for the differential expression of the gene containing, or close to the selected SNP. The output for the rs4680 differential gene expression. A search result for rs4680 SNP is presented as an example. By clicking on the names under the “Column Description” title, the user can see a box, explaining the column or abbreviations that are used in that column in the results section. “Assay” (red box) is selected as an example here.

**Figure S5.**
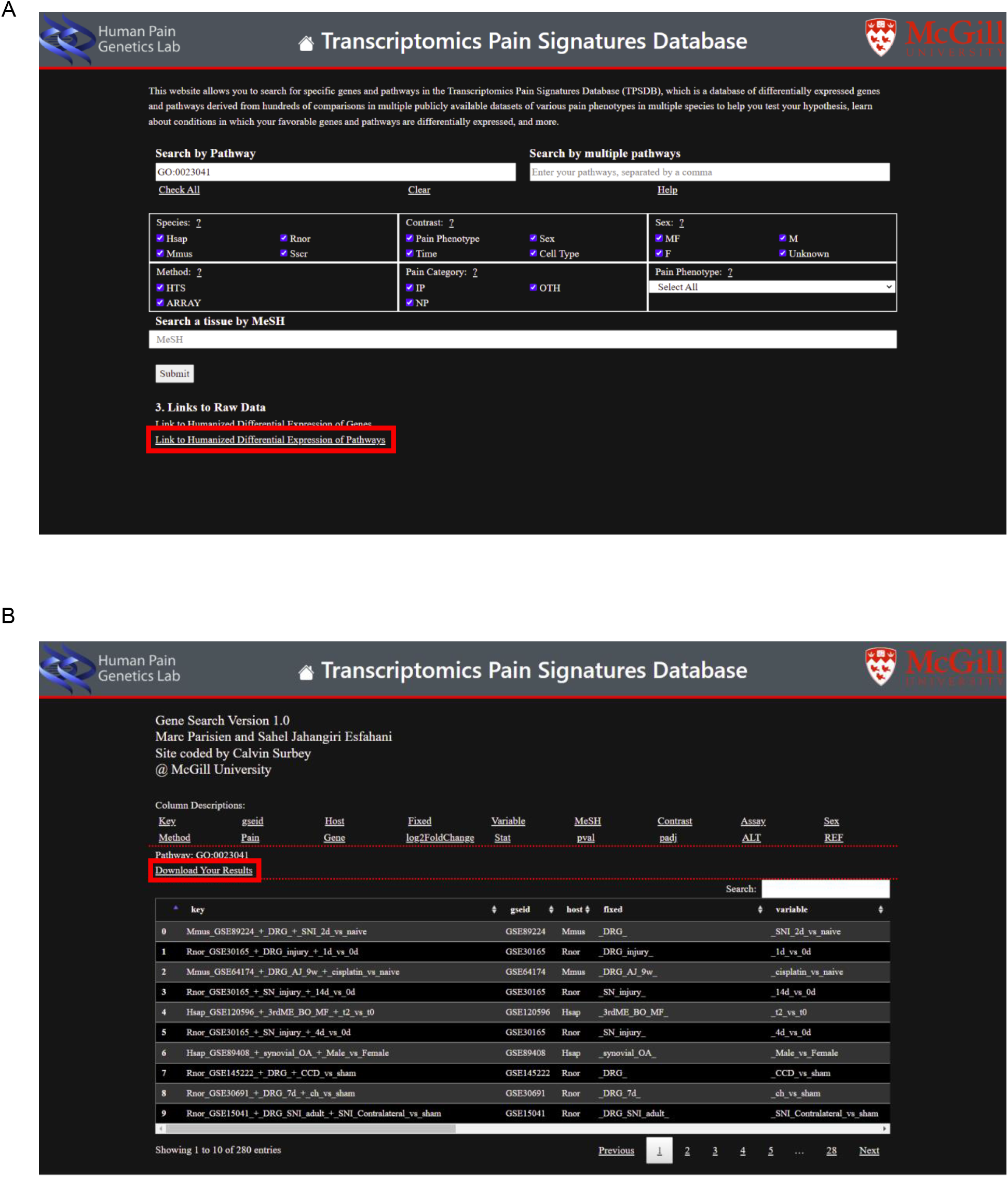
TPSDB website. **(A)** Pathway search page. As an example, the search for the “GO:0023041” pathway in all sexes, species, and pain states is shown. All the results of differentially expressed pathway analysis available in the database, are downloadable by clicking on “Link to Humanized Differential Expression of Pathways” (red box). **(B)** The output for the differential pathway expression. A search result for the “GO:0023041” pathway is presented as an example. The results’ excel file for any specific search is downloadable by clicking on the “Download Your Results” (red box) section.

**Figure S6.**
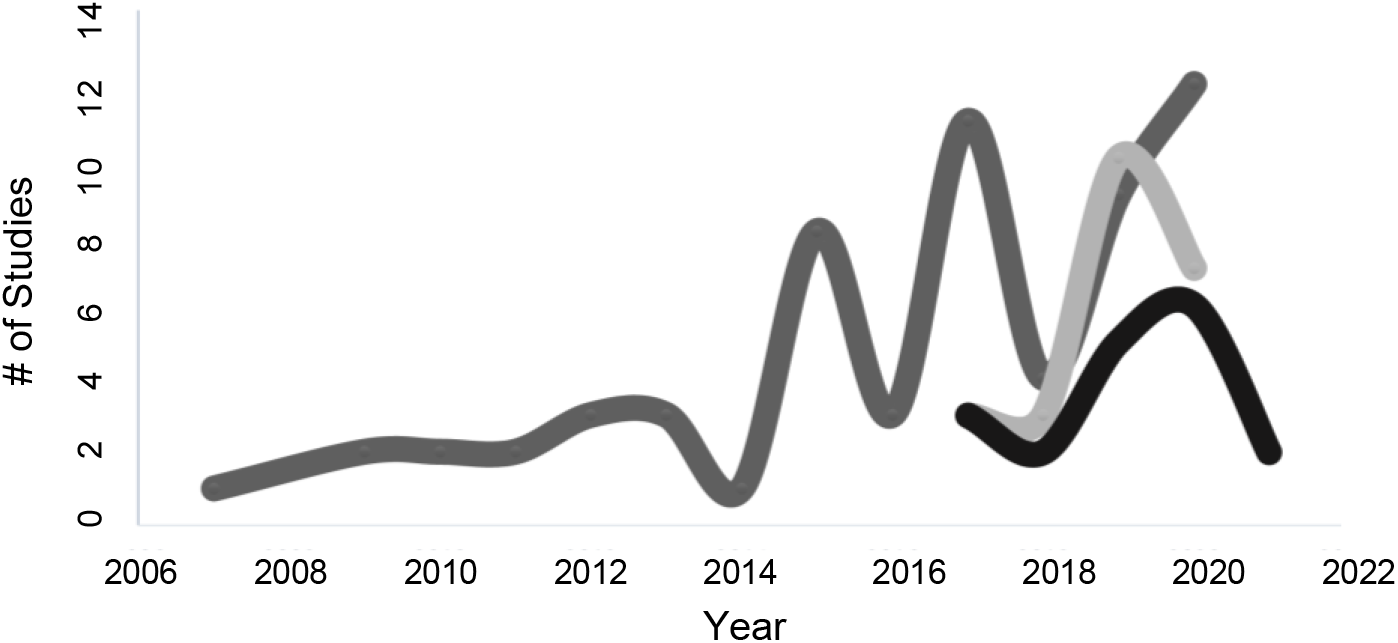
Usage of each sex in pain studies over time. The y-axis shows the number of studies in that year, while the x-axis represents the year. Dark gray represents studies that are done in male. Light gray represents studies that are done in female. Black represents studies that are done in both male and female.

**Figure S7.**
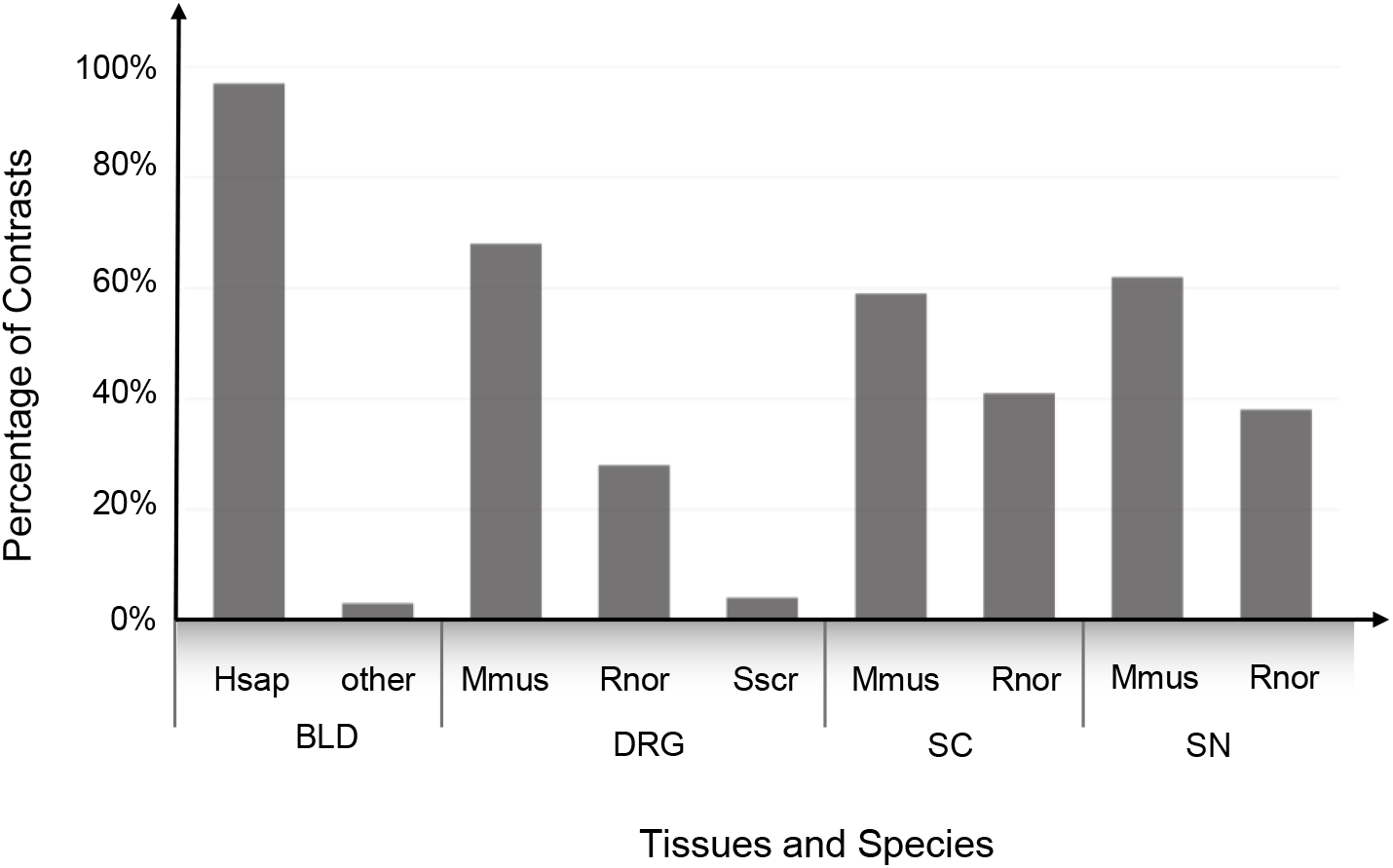
Tissue and specie-wise count of the TPSDB contrasts. The proportion of the contrasts per organism for each tissue is demonstrated in the bar chart. Hsap; Homo Sapiense, Mmus; Mus Musculus, Rnor; Rattus norvegicus, Sscr; Sus scrofa, BLD; blood, DRG; dorsal root ganglion, SC; spinal cord, SN; sciatic nerve. X-axis represents tissues and species. Y-axis represents the percentage of the contrasts.

**Figure S8.**
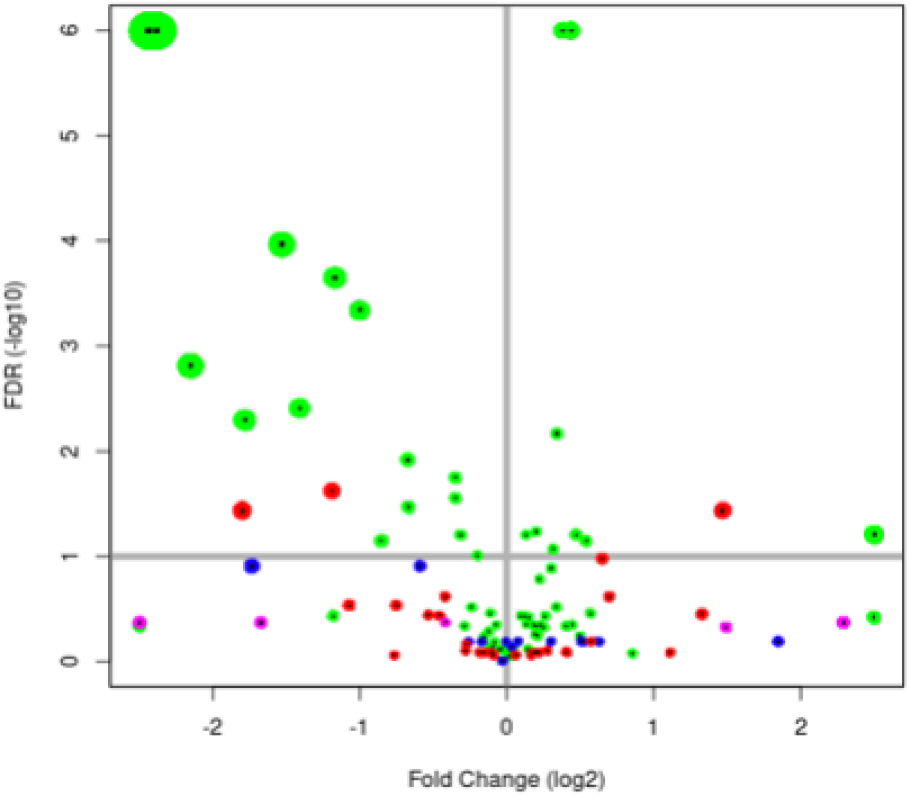
Differential expression of the TRPV1 gene. The TRPV1 gene was found differentially expressed in 28 contrasts at the FDR 10% level, of which 25 contrasts were in DRG, while the TRPV1 gene expression was found in 67+51 contrasts, of which 67 were in the DRG. Tissues are: DRG (green), spinal cord (blue), blood (red), and sciatic nerve (magenta). Each dot is a contrast.

**Figure S9.**
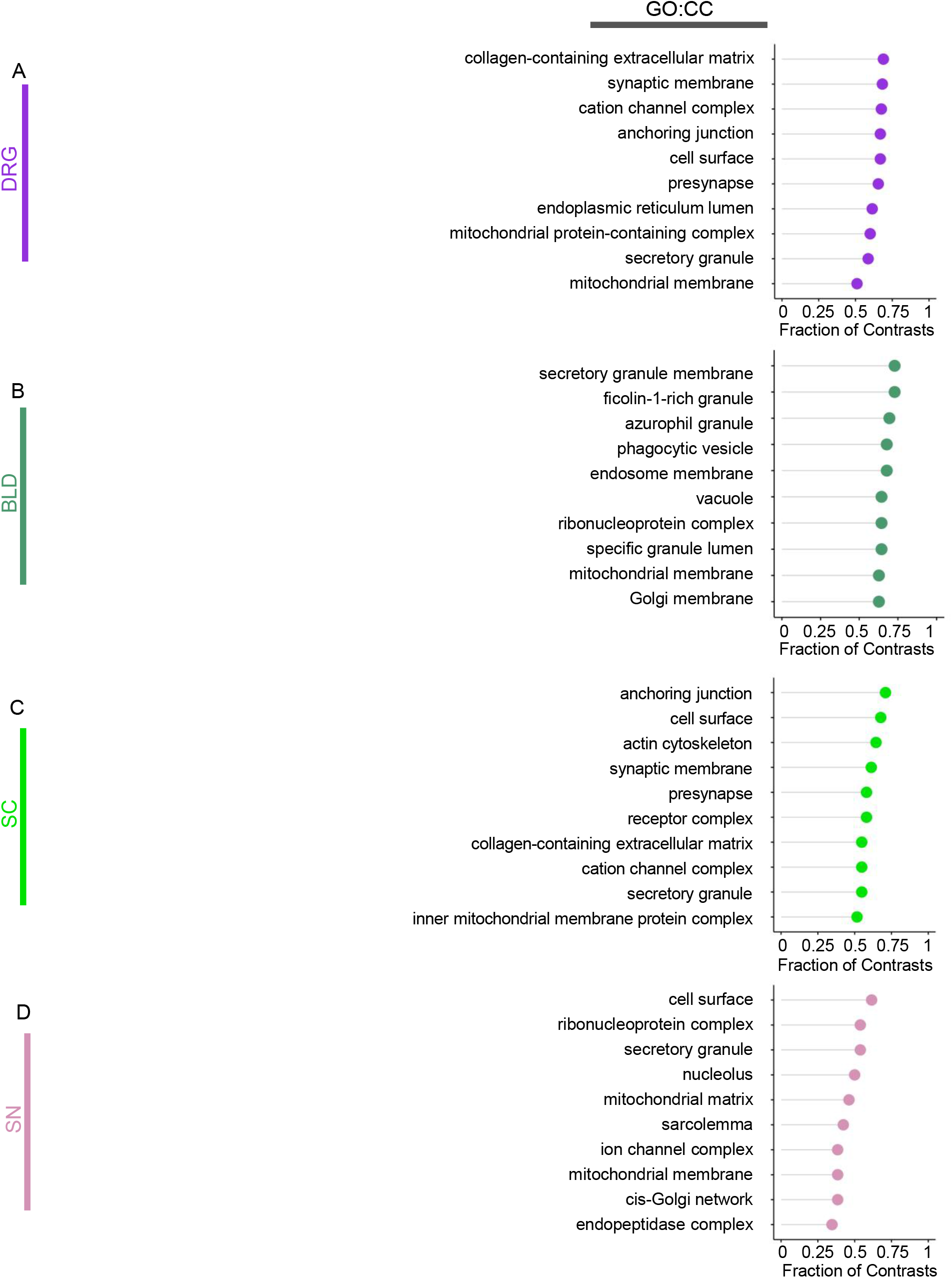
Pathway-level statistics from the TPSDB database. Top ten cellular components (GO:CC) most significantly enriched per tissue. Plots track the proportion of contrasts per tissue where each pathway was most represented in enrichment analysis. (A) dorsal root ganglia (DRG); (B) whole blood (BLD); (C) spinal cord (SC); and (D) sciatic nerve (SN).

**Figure S10.**
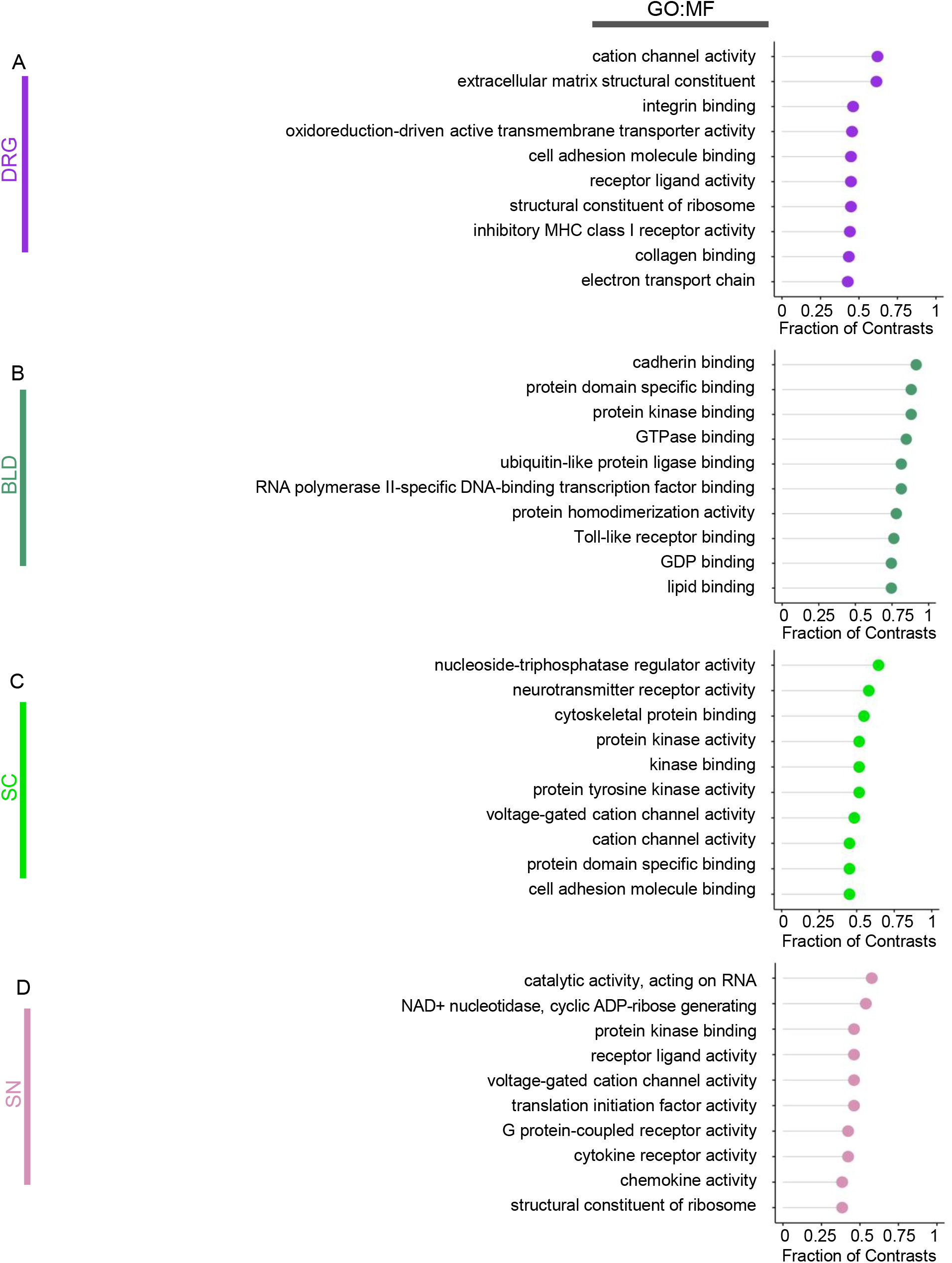
Pathway-level statistics from the TPSDB database. Top ten molecular function (GO:MF) most significantly enriched per tissue. Plots track the proportion of contrasts per tissue where each pathway was most represented in enrichment analysis. (A) dorsal root ganglia (DRG); (B) whole blood (BLD); (C) spinal cord (SC); and (D) sciatic nerve (SN).

**Figure S11.**
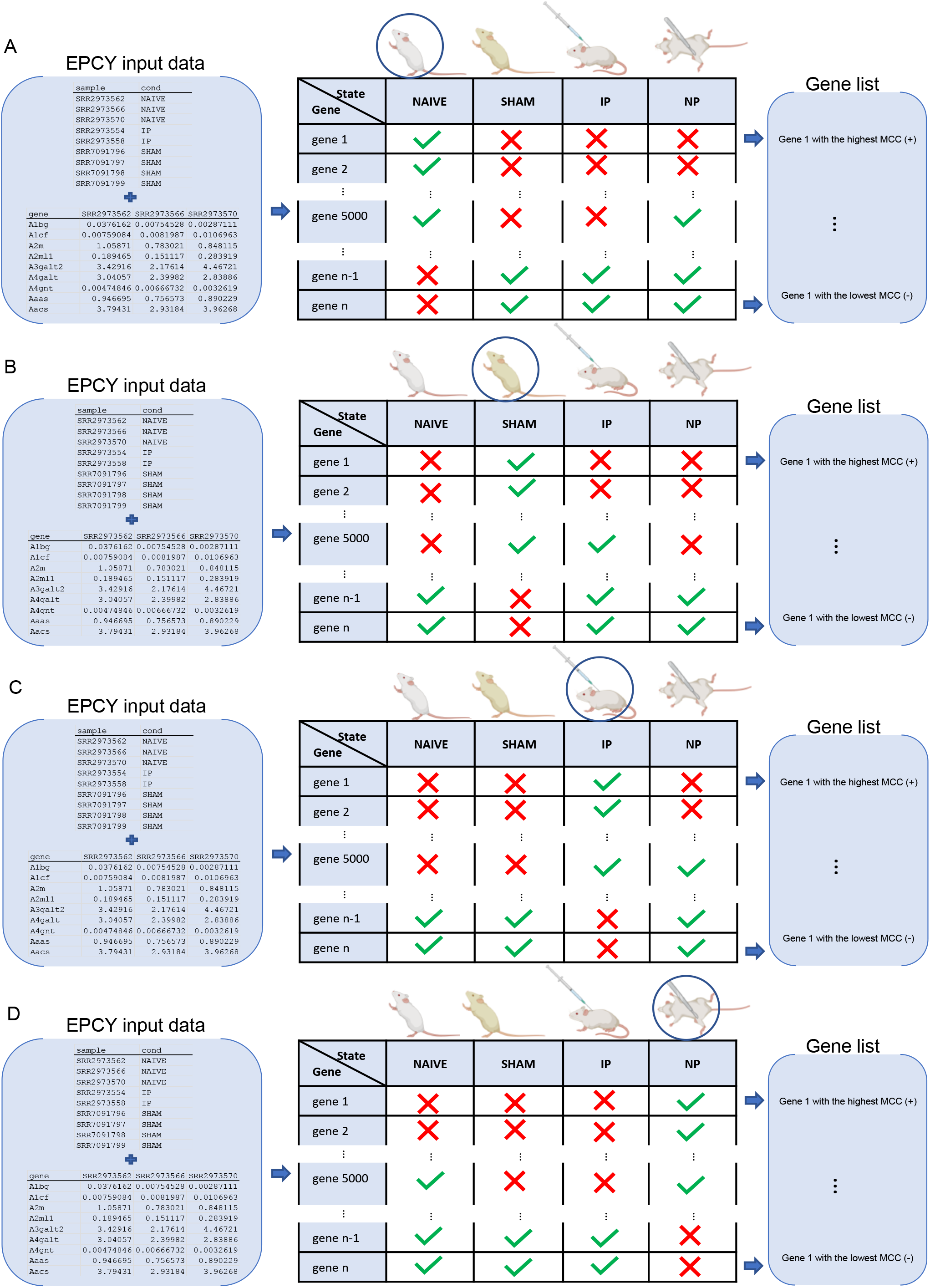
Groups analysis by EPCY approach. EPCY uses input data and ranks each gene based on the direction of correlation with each pain state. The highest matthews correlation coefficient (MCC) in the gene list reflects the highest positive correlation with the pain state and the lowest MCC reflects the negative correlation with the pain state. (A) Predicting NAIVE pain state among other states. (B) Predicting SHAM pain state among other states. (C) Predicting inflammatory pain (IP) among other states. (D) Predicting neuropathic pain (NP) among other states.

