## supplementary tables description for "The Transcriptomics Pain Signature Database"

**Supplementary Table 1. Pain Gene Collection.** Eleven sources were used to create the collection of genes previously implicated in pain. Description sheet contains all the references and the number of genes that are added to the “Pain Gene Collection” from them; (A) Pain Gene Database; (B) Algynomics Pain Res Panel; (C) Human Pain Genetics Database; (D) Pain Genes 2008; (E) Pain Genes 2012; (F) Pain Research Forum; (G) Gene Ontology "pain"; (H) geneRIFs; (I) GeneCards; (J) Comp Toxicogenomics DB; (K) Harmonizome "pain" @ CTD; (L) Final list, containing 2254 genes of “Pain Gene Collection”.

**Supplementary Table 2. Summary of contrasts of the pain states for DEG in TPDB.** (A) Contrasts and their characteristics. (B) Pain assays.

**Supplementary Table 3. Gene-level statistics from the TPSDB.** Columns are: the official gene symbol, ‘gene’; the number of times the gene was significant in all contrasts, ‘nb_signif’; the total number of contrasts where a gene was measured, ‘nb_total’; the fraction of significant occurrences, ‘fraction’. Threshold of significance was set up for FDR 10%. Tissues are: (A) dorsal root ganglia, DRG; (B) whole blood, BLD; (C) spinal cord, SC; (D) sciatic nerve, SN.

**Supplementary Table 4. Enrichment of “popular pain genes” in TPSDB.** Hypergeometric test for “Popular Pain Genes”^22^ was done using differential gene expression results and enrichment scores of these genes. (A) Description of the analysis. (B) The most significant tissue per gene. (C) The results for all of the “popular pain genes”.

**Supplementary Table 5. Pathway-level statistics from the TPSDB.** Statistical significance counts for each pathway by tissues. A pathway was declared significant in a contrast if its FDR-adjusted P-value was equal to or lower than 0.1 (FDR 10%). Pathways from Gene Ontology’s biological processes (GOBP), cellular components (GOCC), and molecular functions (GOMF). Columns are: a Boolean to keep (or not) the pathway, ‘keep’; the pathway’s depth in the GO tree, ‘depth’; the pathway’s height in the GO tree, ‘height’; the proportion of new genes in the pathway compared to those already chosen in the kept pathways, ‘novelty’; the number of genes in the pathway, ‘size’; the pathway's identifier, ‘goid’; the pathway's short description, ‘desc’; the number of times the pathway was significant in all contrasts, ‘nb_signif’; the total number of contrasts, ‘nb_total’; the fraction of significant occurrences, ‘fraction’. A pathway is ‘kept’ if: has minimum depth of 2, maximum height of 5, minimum novelty of 0.5. Tissues are (A) dorsal root ganglia, DRG; (B) whole blood, BLD; (C) spinal cord, SC; (D) sciatic nerve, SN.
